# Neurotrophin receptor Ntrk2b function in the maintenance of dopamine and serotonin neurons in zebrafish

**DOI:** 10.1101/136416

**Authors:** Madhusmita Priyadarshini Sahu, Ceren Pajanoja, Stanislav Rozov, Pertti Panula, Eero Castrén

**Affiliations:** Neuroscience Center, Helsinki Institute of Life Science HiLife, University of Helsinki, 00290 Helsinki, Finland; Institute of Biomedicine/Anatomy, University of Helsinki, 00290 Helsinki, Finland

## Abstract

Brain-derived neurotrophic factor (BDNF), together with its cognate receptor tyrosine kinase B (TrkB), plays an essential role in the development and plasticity of the brain and is widely implicated in psychiatric diseases (Autry and Monteggia, 2012). Due to the highly conserved evolutionary lineage of neurotrophins and their receptors in vertebrates, the zebrafish is a well-suited model for this study. The TrkB receptor, also known as NTRK2, has two forms in zebrafish, Ntrk2a and Ntrk2b. The spatio-temporal expression pattern of *bdnf* and *ntrk2b* in zebrafish was studied using *in situ* hybridization. The complementary expression pattern of *ntrk2b* to *bdnf* suggests that *ntrk2b* is the key receptor, unlike its duplicate isoform *ntrk2a*. Two reverse genetics strategies, morpholino oligonucleotides (MO) and the TILLING mutant, were applied in this study. The loss or complete deletion of *ntrk2b* had no major effect on the viability, gross phenotype, or swimming behavior of zebrafish. A specific subset of the dopaminergic and serotonergic neuronal population was affected in the morphants and mutants. Downstream signaling transcripts such as *bdnf, serta*, *th2*, and *tph2* were downregulated and could be rescued by overexpression of the full-length *ntrk2b* mRNA in the morphants. Pharmacological intervention with a tyrosine kinase inhibitor, K252a, resulted in similar phenotypes. Overall, our results reveal a specific effect of *ntrk2b* on the two crucial aminergic systems involved in psychiatric disorders and provide an essential tool to study neurotrophin function in modulating neuronal plasticity in the central nervous system.

**Significance Statement:** Brain-derived neurotrophic factor (BDNF) and its high-affinity receptor, tyrosine kinase (TrkB/NTRK2), play a major role in regulating the development and plasticity of neural circuits. Additionally, BDNF/TrkB signaling is involved in psychiatric disorders and antidepressant responses. This study presents the complementary gene expression pattern of TrkB and BDNF in zebrafish during the early larval stage and in the adult brain. Our results consistently indicate that BDNF/TrkB signaling has a significant role in the development and maintenance of dopaminergic and serotonergic neuronal populations. Therefore, the *ntrk2b*-deficient zebrafish model is well suited to studying psychiatric disorders attributed to a dysfunctional monoaminergic system, and could potentially be a valuable model for small molecule drug screening.

## Introduction

Neurotrophins (NTs) are a family of receptors known to play critical roles in nervous system development, maintenance, and synaptic plasticity (Huang and Reichardt, 2001). The neurotrophin family includes nerve growth factor (NGF), brain-derived neurotrophic factor (BDNF), neurotrophin-3 (NT-3), neurotrophin-4/5 (NT-4/5), neurotrophin-6 (NT-6), and neurotrophin-7 (NT-7). They bind with high affinity to Trk neurotrophin receptors (NTRK2), transmembrane tyrosine kinase proteins TrkA, TrkB, and TrkC (Huang and Reichardt, 2001). Trk receptors mediate the trophic properties of all neurotrophins. Both NTs and Trk receptors are phylogenetically highly conserved amongst vertebrates (Huynh and Heinrich, 2001).

Neurotrophin BDNF binds with high affinity to TrkB and plays a prevalent role in neuronal plasticity. Complete knockouts for studying BDNF or TrkB deficiency have been unsuccessful in rodent models due to developmental abnormalities and respiratory failure leading to postnatal lethality (Erickson et al., 1996). Most studies have been carried out on conditional knockout rodents using different gene specific promoters (Li et al., 2008). BDNF/TrkB signalling has been linked with both the pathophysiology of depression and the mode of action of antidepressants (Castren and Hen, 2013; Castren and Rantamaki, 2010). The neurotransmitter serotonin has been associated with depression and is a major target for antidepressant treatment, especially the selective serotonin reuptake inhibitors (SSRIs) (Martinowich and Lu, 2008; Gorman and Kent, 1999). BDNF/TrkB molecules are co-localized in neurons of the raphe nucleus, the serotonin-producing region in the brain. They both can co-regulate each other (Mattson et al., 2004; Madhav et al., 2001; Mamounas et al., 2000). The bidirectional effect of these two major systems needs to be further explored to be able to understand the delay in the mode of action of antidepressants in human patients.

BDNF/TrkB has primary impact on the reward circuitry governed by the dopaminergic circuit.(Autry and Monteggia, 2012; Wook Koo et al., 2016). Depressive-like behavior has been associated with the mesolimbic dopaminergic pathway (Wook Koo et al., 2016). The depression-like effect produced by BDNF in the mesolimbic circuit is contradictory to its antidepressant-like effect in the hippocampus (Nestler and Carlezon, 2006; Berton et al., 2006). The interaction between BDNF/TrkB and dopamine signaling in the brain is important in understanding the effect of stress on depression, and also for addictive behavior in the functional reorganization of neuronal networks in psychiatric disorders.

The zebrafish is an extensively used vertebrate model due to its high fecundity, short generation time, and similar neuroanatomy to the mammalian brain (Panula et al., 2010). With the advancements in knockdown and knockout techniques in zebrafish, it is increasingly used as an animal model for human biology and disease. There are five trk receptors in the zebrafish genome, Trka/Ntrk1, Trkb1/Ntrk2a, Trkb2/Ntrk2b, Trkc1/Ntrk3a, and Trkc2/Ntrk3b (Martin et al., 1995). The transcript *ntrk2b* has been confirmed to be similar to mammalian TrkB, as *ntrk2a* has not been robustly detected (Gasanov et al., 2015). BDNF knockdown in zebrafish results in severe phenotypic abnormalities (Diekmann et al., 2009). Applying a reverse genetics strategy, we aimed to demonstrate the effects of Ntrk2b loss and its consequences for the aminergic systems in the larval and adult zebrafish brain.

## Materials and methods

### Zebrafish strain and maintenance

The zebrafish strain used in the experiments was the wildtype Turku strain. It has been maintained in our facility for more than a decade (Kaslin et al., 2004; Chen et al., 2009; Sallinen et al., 2010). Animals were raised at 28 °C and staged as described earlier (Kimmel et al., 1995). The mutant for Ntrk2b (Sa13660) was obtained from the zebrafish mutation project (Wellcome Trust Sanger Institute) and backcrossed with the wildtype Turku strain at least twice. All experimental procedures were performed in accordance with institutional animal welfare guidelines and were approved by the Office of the Regional Government of Southern Finland in agreement with the ethical guidelines of the European convention (ESAVI/10300/04.10.07/2016).

### Full-length *ntrk2b* construct

The full-length *ntrk2b* was synthesized using the following primers:

F:5’-GGATCCCGCTAGACCTGCTATGACCG-3’,

R: 5’-GGATGTCCAGGTACACAGGCCCTAGG-3’.

The PCR cycling parameters were 94 °C for 2 min and 40 cycles of 94 °C for 30 sec, 58 °C for 30 sec, 72 °C for 2 min, followed by an extension at 72 °C for 10 min. The PCR fragment was cloned into PGEMT-easy vector systems according to the manufacturer’s instructions (Promega, Madison, WI) and was verified by sequencing. The BDNF clone was obtained from Prof. Pertti panula and was sequence verified.

### Morpholino oligonucleotide (MO) design and mRNA rescue injections

The antisense morpholino oligonucleotides were ordered from GeneTools LLC. The translation start site was targeted for designing the Ntrk2b MO (5´-TTCCACGAACCCCTGCGGTCATAGC). The working concentration was determined with a dose-dependent assay. A standard control MO was used as a control (Priyadarshini et al., 2013). The full-length *ntrk2b* sequence was artificially synthesized and the codon optimized for the mRNA rescue experiments (Geneart^TM^, Life Technologies, Carlsbad, USA). Additional restriction sites were inserted to subclone it to the pMC vector containing untranslated repeats and polyadenylation, which can enhance mRNA stability and translation efficiency. The pMC vector was kindly provided by Dr Thomas Czerny. The pMC vector containing the full-length *ntrk2b* insert was linearized with NotI, and capped full-length transcripts were generated with an mMESSAGE mMACHINE kit (Ambion, Austin, Tx) using T7 polymerase. For the mRNA rescue, 1 ng of *ntrk2b* mRNA was co-injected with 4 ng Ntrk2b MO at the one-cell stage.

### In situ hybridization

Fish at 3 dpf were fixed in 4% PFA in phosphate buffered saline (PBS) o/n at +4 °C. The fixative was changed the next day to 100% methanol and the samples were stored at -20 °C until further use. The adult brains were dissected and fixed with 4% PFA o/n at +4 °C. They were embedded in cryo-embedding mix until solidified. The solidified brains were sectioned at 14 μm on Superfrost slides and kept at -80 °C until further use. The *in situ* hybridization method applied has been described earlier (Thisse and Thisse, 2008). The probes were synthesized from the clones *serta* (a kind gift from K. Shirabe), *th1, tph1a, tph2, ntrk2b*, and *bdnf*. The digoxigenin (DIG)-labeled probes were generated with the DIG RNA labeling kit according to the manufacturer’s instructions (Roche, Mannheim, Germany). The samples were mounted in 80% glycerol and brightfield images were taken using a Leica DM IRB inverted microscope with a DFC 480 charge-coupled device camera, and z-stacks were processed with Leica Application Suite software and CorelDRAW 12 software (Synex, Brooklyn, USA).

### Western blotting

The adult mutant zebrafish brains and MO-injected fish at 2 dpf were collected in standard RIPA lysis buffer (3M Tris HCl pH 8, 5M NaCl, 0.5M NaF, NP-40, glycerol, and protease inhibitor cocktail tablets). Homogenized fish were centrifuged at 13 000 rpm for 15 minutes at +4 °C. The supernatant was collected and protein estimation was performed using the DC assay kit (Biorad, USA). An equal amount of protein was loaded onto an SDS-PAGE gel after heat denaturation. The proteins were blotted on a PVDF membrane following incubation with the antibodies. The primary antibodies used were TrkB (SC-11, Santacruz), pTrkB (a kind gift from Dr Moses Chao), actin (A4700, Sigma-Aldrich), and GAPDH (Cell Signalling Technology). The secondary antibodies were hrp-conjugated rabbit for trk and GAPDH and mouse for actin (BioRad, USA). The membrane was developed by chemiluminescence using the Pierce ECL kit (Thermo scientific, USA) followed by imaging using a Fuji LAS-3000 Camera (Tamro Medlabs, Vantaa, Finland).

### Quantitative real-time PCR

Total RNA was isolated from pooled whole zebrafish larvae using a PureLink(r) RNA Mini Kit (Thermo Fisher). There were 30 animals per group in triplicates. The RNA was reverse transcribed using the SuperScript IV reverse transcriptase (Invitrogen) primed with oligo (dT) primers according to the manufacturer’s instructions. The primers used were as follows:

zfbactin-1F-CGAGCAGGAGATGGGAACC, zfbactin-1R-CAACGGAAACGCTCATTGC

zfth1F-GACGGAAGATGATCGGAGACA, zfth1R-CCGCCATGTTCCGATTTCT

zfth2 F-CTCCAGAAGAGAATGCCACATG,zfth2R-ACGTTCACTCTCCAGCTGAGTG

zftph2F-GCAAATACTGGGCTCGGAGA, zftph2R-GAGCATGGAGGATGCAAGGT

zftph1bF-GCCTGTTGCTGGCTATCTGT, zftph1bR-CTCATGACAGGTGTCCGGTTC

zftph1a F-TCTACACACCTGAGCCAGAC, zftph1a R-CCCTTCCTGCTTACAGAGCC

zfbdnf F-CTCGAAGGACGTTGACCTGT, zfbdnf R-CGGCATCCAGGTAGTTTTTG

The results were analyzed using GraphPad Prism software (GraphPad Software, Inc.CA, USA).

### Immunostaining

All the fish embryos were fixed at 5 dpf with 4% PFA overnight at +4 °C. The samples were processed as published earlier (Kaslin et al., 2004;(Sallinen et al., 2010). The primary antibodies used were anti-tyrosine hydroxylase monoclonal mouse antibody (MAB318, 1:1000; Chemicon) and anti-serotonin rabbit antibody (S5545, 1:1000; Sigma, St. Louis, MO). The secondary antibodies were Alexa Fluor® 488 or 568 goat anti-mouse or anti-rabbit IgG (1:1000; Invitrogen, Eugene, OR). Immunofluorescence samples were mounted in 80% glycerol and examined under a Leica TCS SP2 AOBS confocal microscope. For excitation, an argon laser (488 nm) was used. Emission was detected at 500–550 nm, as described earlier (Kaslin et al., 2004). Stacks of images taken at 1.2-µm intervals were compiled, and final images were produced with Leica Confocal Software using the maximum intensity projection algorithm.

### Amine levels and behavioral analysis

The levels of serotonin and dopamine in the whole larval fish at 3 dpf were analyzed by high pressure liquid chromatography (HPLC). We used 30 larval fish per sample, which were lysed in 150 µl of 2% perchloric acid by sonication. After centrifugation, 10 μl of supernatant was analyzed for the monoamine concentration by HPLC. Three individual groups per treatment condition were measured as a blinded experiment. Larval zebrafish at 5 dpf were used for a locomotor assay, as has previously been described (Sallinen et al., 2010)

### Drugs

The tyrosine kinase inhibitor K252a was used in this study (Calbiochem #420298, USA). The dosages used in our experiments were 100 nM and 200 nM for the experimental group, and the control was treated with 1% DMSO. The zebrafish were treated with the compound for 24 hours, after which they were immediately sacrificed and the brains were dissected for further processing.

## Results

### Transcript for *ntrk2b* and *bdnf* expression in zebrafish

The spatiotemporal expression of the two transcripts was examined at 3 dpf and in the adult brain sections by whole-mount *in situ* hybridization. The full-length *ntrk2b* mRNA was widely expressed in the brain and spinal cord at 3 dpf (Fig. 1A–B). The expression was also visible in the retinal region of the eye. No expression was observed in the sense probe (Fig. 1C). To characterize the expression in the adult brain, sagittal sections of one-year-old adult fish brain were used for *in situ* hybridization. Expression of *ntrk2b* was observed in the dorsal telencephalon, the pallium, in the parvocellular pre-optic nucleus, the posterior tuberculum, radial glial cells lining the mesencephalic ventricle, the cerebellum, the hypothalamus, and a dispersed staining pattern in the medulla oblongata (Fig. 1D).

**Fig 1.**
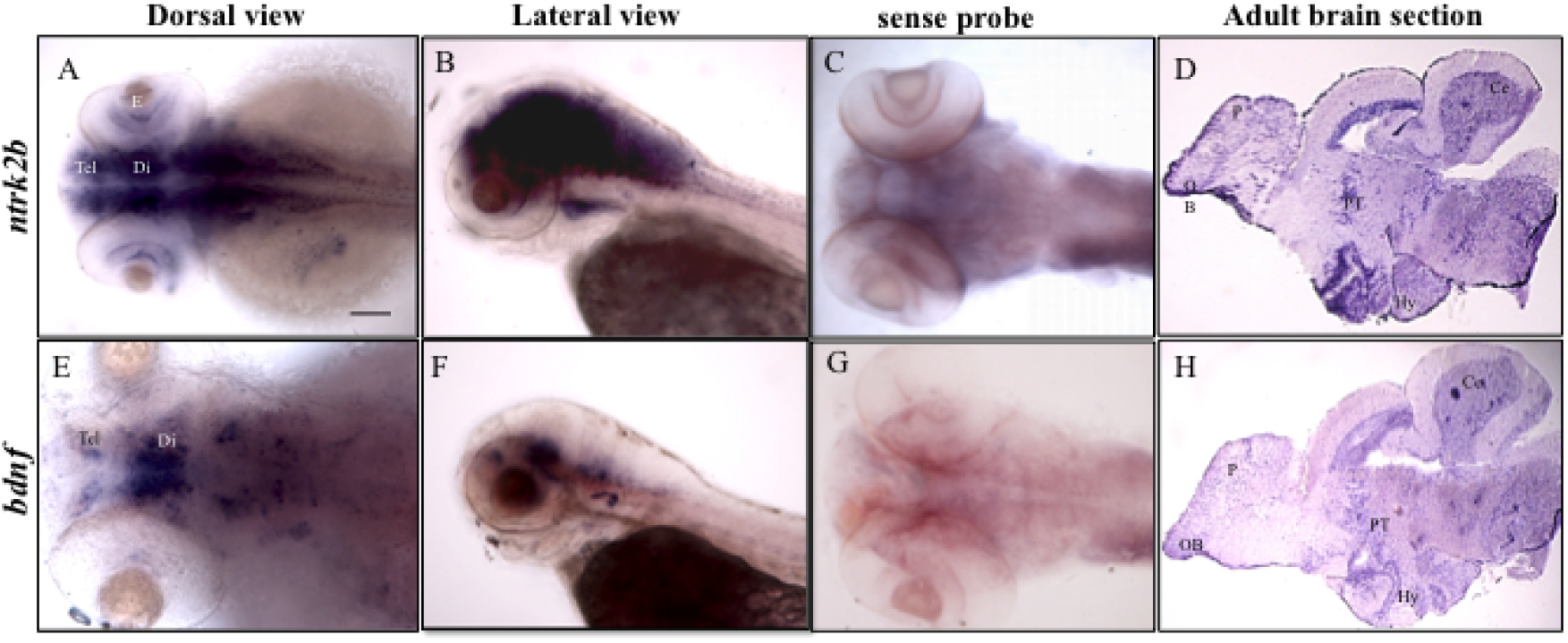
Comparative expression patterns of ntrk2b and bdnf transcripts. a–b: *ntrk2b* antisense expression revealed by *in situ* hybridization at 3 dpf, anterior to the left. c: Sense probe for *ntrk2b* at 3 dpf. d: *ntrk2b* expression in a 1-year-old adult brain section, anterior to the left and dorsal at the top. e–f: *bdnf* expression in a larval brain, anterior to the left. g: Sense probe for *bdnf* at 3 dpf. h: *bdnf* expression in a 1-year-old adult brain section, anterior to the left and dorsal at the top. Tel - telencephalon, Di - diencephalon, E - eye, Hy - hypothalamus, Ce - cerebellum, PT - posterior tuberculum, P - pallium, OB - olfactory bulb. Scale bar = 100 µm.

The *bdnf* transcript at 3 dpf had a rather restricted expression pattern. It was visible in the telencephalon, the pre-optic region in the diencephalon, and in the rhombomeres (Fig. 1E–F). The sense probe for *bdnf* served as the negative control (Fig. 1G). Complementary expression for *bdnf* was detected in the adult brain in similarly represented regions as *ntrk2b* (Fig. 1H).

### Both translation blocking morpholino oligonucleotides and a point mutation attenuated *ntrk2b* function

The function of Ntrk2b in zebrafish was investigated by applying two different reverse genetics methods, morpholino oligonucleotide (MO)-based translation inhibition and a TILLING mutant. The Ntrk2b TILLING mutant was obtained from the zebrafish mutation project at the Wellcome Trust Sanger Institute. The mutant has a point mutation (T > A) at exon 17, which results in a non-functional Ntrk2b protein. These mutants were bred and genotypes were grouped after sequence verification. These mutants have been outcrossed to eliminate the possible effects of non-specific mutations. The Ntrk2b mutant had no gross phenotype as compared to its wildtype littermate (Fig. 2A). Using a translation-blocking morpholino, we observed no morphological differences between the standard control-injected and ntrk2b MO-injected embryos at 1 dpf and 5 dpf (Fig. 2B). To assess the knockdown efficiency, the embryos were injected with different doses (2 ng, 4 ng, and 6 ng) of the ntrk2b MO and collected at 2 dpf. The deletion efficiency was confirmed by Western blotting using a polyclonal antibody against TrkB (Fig. 2C). A faint band was observed at 2 ng, while no bands were detected at the 4 ng dose. Successful knockdown for Ntrk2b was observed at the 4 ng dose. Actin was used as the loading control for all the samples. Adult brains were isolated from wildtype and heterozygous mutant fish and subjected to Western blot analysis. The heterozygous mutants had a partial loss of TrkB protein (Fig. 2D). GAPDH was used as the loading control. The complete or partial loss of Ntrk2b did not affect the swimming behavior of the mutants or the morphants (Fig. 2E and Fig. 2F, respectively). The morphants and mutants behaved similarly compared to the respective controls in the quantitative locomotor assay. We quantified the *bdnf* transcripts in these MO-injected animals and found a significant reduction in *bdnf* mRNA levels as compared to the control-injected ones (Fig. 2G). This decreased transcript for *bdnf* could be rescued by *ntrk2b* mRNA injection of the MO-injected animals.

**Fig 2.**
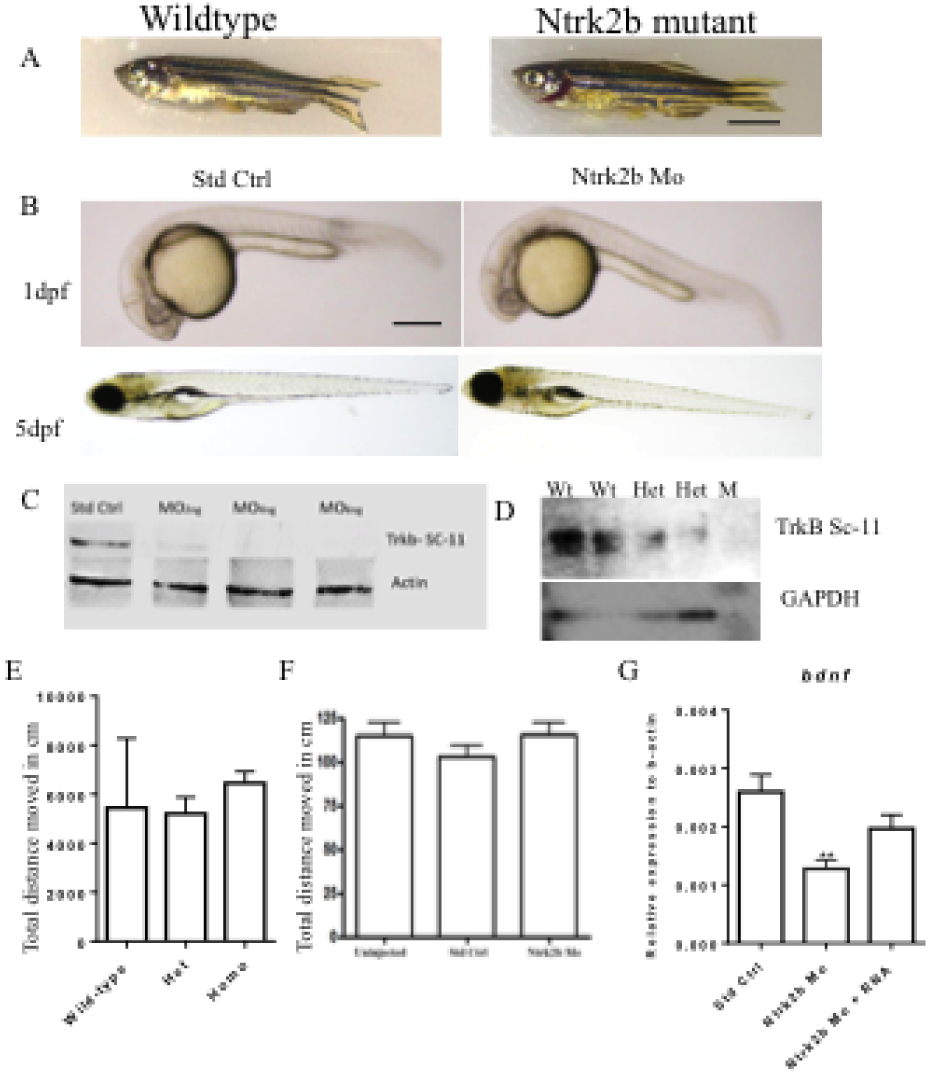
Ntrk2b deletion efficiency with no major effect on morphology and motor function, but the levels of bdnf were reduced. a: Gross phenotypic difference between age-matched wildtype and Ntrk2b mutant fish. Scale bar = 500 mm b: No morphological effect of standard control-injected and morpholino-injected fish at 24 hours after injection and 5 dpf. c: Western blotting with Trk antibody SC-11 to check for knockdown efficiency in the dose-dependent injection for Ntrk2b MO. Actin antibody was used as the loading control. d: Age-matched wildtype and heterozygous adult mutant brain samples verified with Trk antibody SC-11. GAPDH was used as the loading control. e: Quantitative locomotor analysis revealed no significant difference between wildtype, heterozygous, and homozygous mutant fish. f: Motor behavior in the uninjected control, standard injected control and ntrk2b morpholino remained unaltered. g: Q-RT PCR for the bdnf transcript in the ntrk2b morphant. **p =0.0029 one-way ANOVA. Scale bar = 100 µm.

### Effect of *ntrk2b* inhibition on the dopaminergic system

BDNF/TrkB signaling has been associated with dopaminergic signaling in the mesolimbic circuitry in relation to chronic stress (Wook Koo et al., 2016). Transient inactivation of *ntrk2b* by morpholino oligonucleotides reduced the total dopamine levels in the larval zebrafish at 3 dpf, as determined using HPLC (Fig. 3A). There are two complementary forms of tyrosine hydroxylase in the zebrafish: *th1* and *th2*. Quantitative estimation of the transcript levels with real-time PCR revealed significantly reduced levels of both *th1* and *th2* (Fig. 3Ba–b). The decreased transcript levels could be rescued by overexpression of full-length mRNA for Ntrk2b-injected morphants.

**Fig 3.**
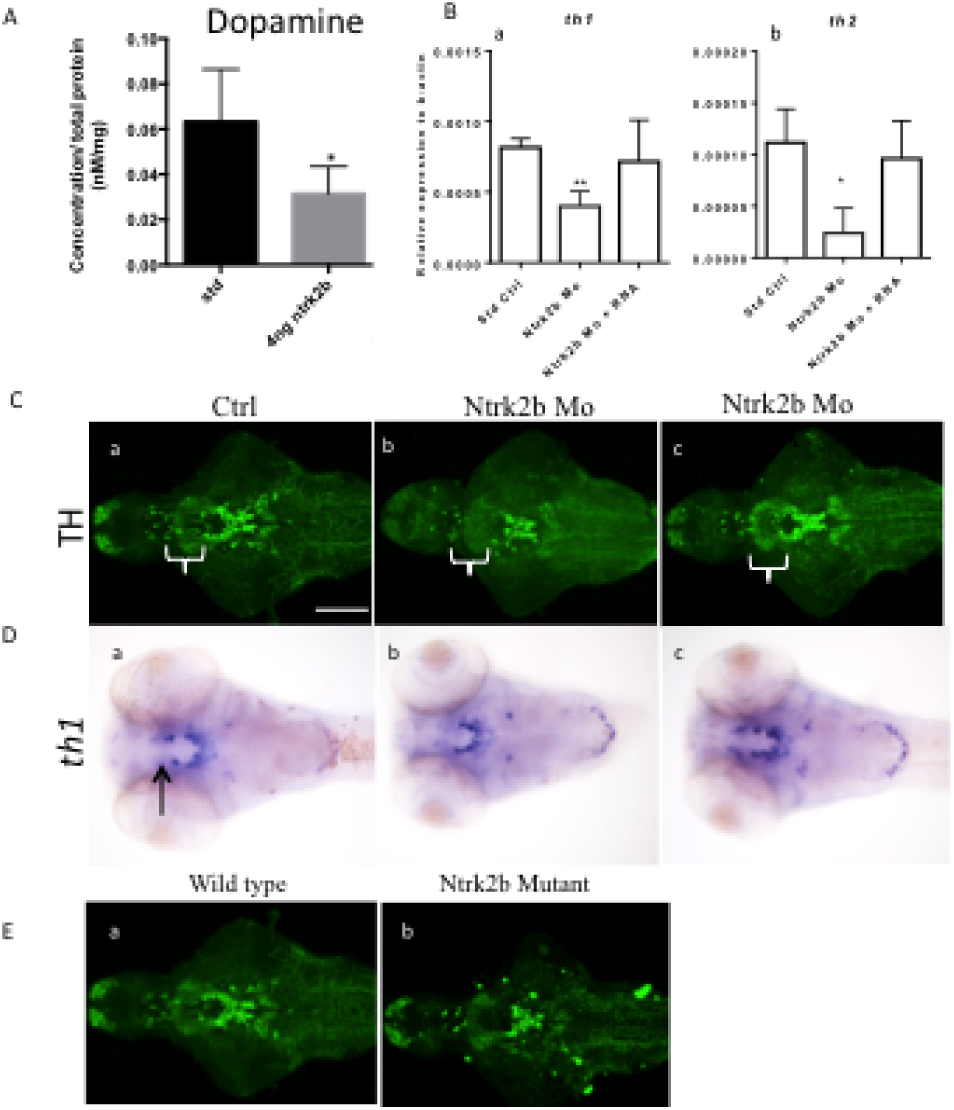
Effect of Ntrk2b deletion on dopamine and its markers. A: HPLC for total dopamine levels in the ntrk2b morphants at 3 dpf revealed them to be reduced compared to controls. B (a–b): Dopamine transcript markers *th1* and *th2* were significantly reduced in the morphant. This effect could be rescued with *ntrk2b* mRNA. C (a–c): TH1 immunohistochemistry revealed a loss of the neuronal population in the diencephalic cluster in the morphant. Ca: The neuronal cell group affected in the control is marked. Cc: The loss of function was restored in mRNA overexpression. D (a–c): ISH for *th1* revealed a similar loss of expression in the diencephalon. E (a–b): The Ntrk2b mutant showed reduced TH immunoreactivity. * p =0.0342, **p=0.0034 one-way ANOVA. Scale bar = 100 µm.

To define the cell populations that could specifically be altered by Ntrk2b inhibition, we applied immunohistochemistry using an antibody for TH1. This antibody has been found to only recognize TH1 neurons (Chen et al., 2009). The neuronal populations expressing the two forms of TH have been well characterized in the zebrafish (Chen et al., 2009; Yamamoto et al., 2010; Rink and Wullimann, 2002). We found that a specific ventral diencephalic population numbered 5,6,11 by Sallinen et al. was reduced in the morphants compared to the controls (Fig. 3C a–c).(Sallinen et al., 2009). This numbering of TH1 cell populations has been followed in this paper, because it comprises all groups, including those that are superimposed on each other in the horizontal representation of cell groups. These specific neurons could be rescued by *ntrk2b* mRNA overexpression in the morphants (Fig. 3C). Using whole-mount *in situ* hybridization (WISH) for the *th1* transcript, a similar loss of the neuronal population was seen at 3 dpf (Fig. 3Da–c). The diencephalic *th1-*positive cells were lost in the morphants and could be rescued by overexpression of *ntrk2b* mRNA. In the ntrk2b mutant, the TH1 immunoreactivity was also assessed to confirm the results. Remarkably, the TH1 neuronal population in the ventral diencephalic cluster and posterior tuberculum was reduced in the homozygous mutants (Fig. 3Ea–b). No major change was observed in the pretectal, olfactory bulb, or subpallial Th1-expressing neurons. These findings were consistent in both the morphants and mutants.

### Effect of *ntrk2b* deletion on the serotonergic system

TrkB is involved in antidepressant-mediated responses, especially to selective serotonin reuptake inhibitors (Castren and Rantamaki, 2010). To elucidate the role of the loss of ntrk2b on the total serotonin levels, we performed HPLC on the morphants. There was a significant reduction in serotonin levels in the morphants as compared to the controls (Fig. 4A). The serotonergic centers in the brain have been mapped in many vertebrates based on the expression of the rate-limiting enzyme tryptophan hydroxylase (tph) and serotonin transporter (Sert) (Lillesaar et al., 2009). The *tph* levels in ntrk2b were quantified by qPCR, and all the three isoforms were analyzed. The transcript levels of the rate-limiting enzymes *tph1a, tph1b*, and *tph2* were quantified (Fig. Ba–c). The transcript *tph2* was significantly reduced (Fig. 4Ba). To study the serotonergic cell groups that were affected, we immunostained the larval brains using a serotonin-specific antibody that has been well characterized in the zebrafish (Chen et al., 2009; Sallinen et al., 2009). We found a significant reduction in 5-HT immunoreactivity in the morphants compared to the controls (Fig. 4Ca–c). The loss of neuronal cells and axonal projection was disrupted in the morphants (Fig. 4Cb). This loss could be rescued by overexpression of ntrk2b mRNA (Fig. 4Cc). To investigate the effects of ntrk2b on serotonin transporter levels, we used *serta,* which shares the highest homology with human and rodent Sert (Wang et al., 2006). The expression of *serta* in the raphe and ventral posterior tuberculum overlaps with the location of serotonin immunoreactivity and expression of *tph2* in the zebrafish (Wang et al., 2006). The *serta* expression at 3 dpf was visible in the raphe nuclei and ventral posterior tuberculum (Fig. 4Da–c). In the ntrk2b MO, *serta* expression was abolished in the ventral posterior tuberculum (Fig. 4Db). Few cells in the raphe nuclei were visible in the morphants. Overexpression with ntrk2b mRNA rescued both of the cell groups (Fig. 4Dc).

**Fig 4.**
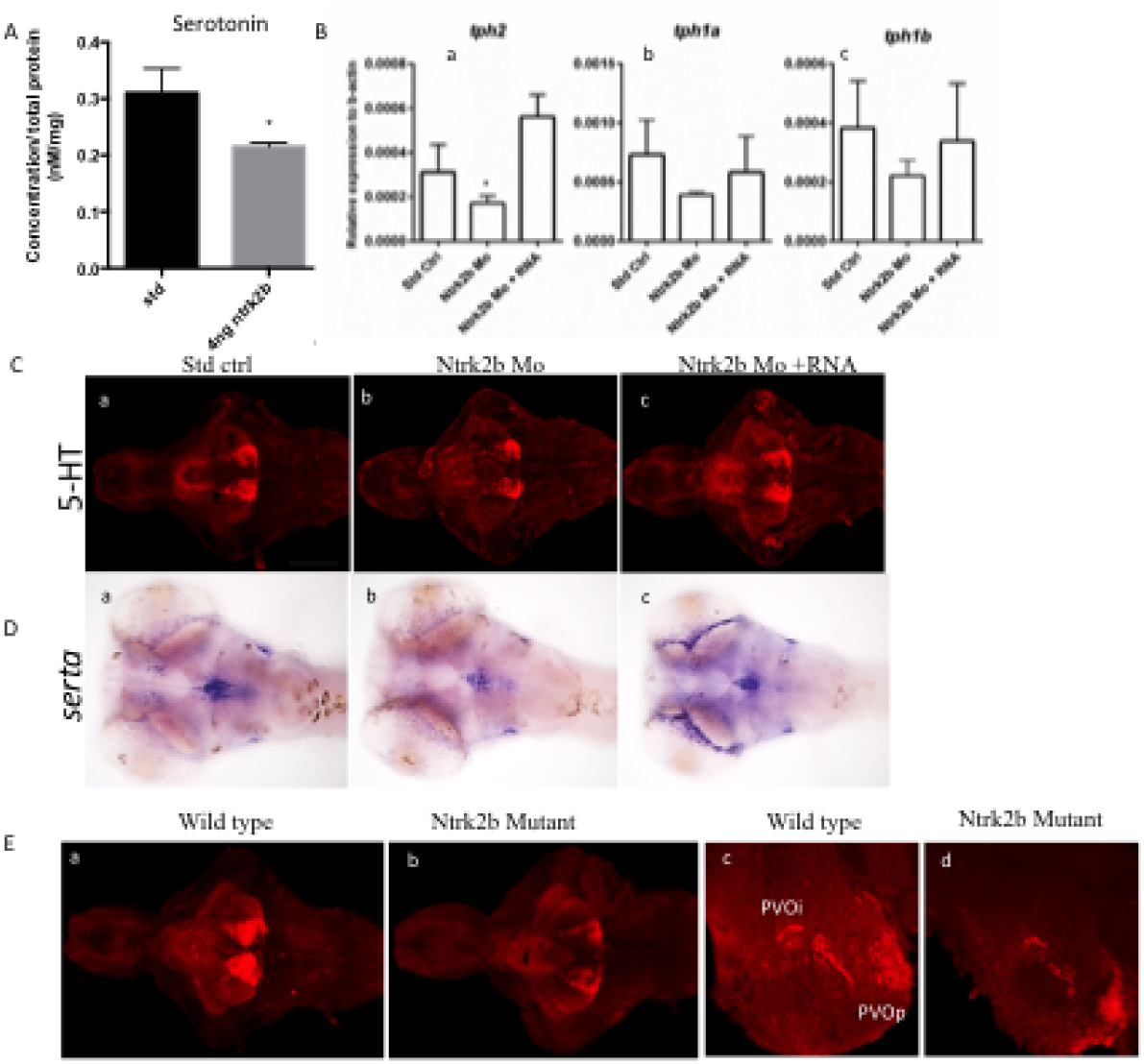
Loss of Ntrk2b effects on serotonin levels and its markers. A: Total serotonin levels were reduced in the morphants. B: The three TPH transcripts were analyzed in the morphants by QRT-PCR: Ba - tph2, Bb - tph1a, Bc - tph1b. Ba: Only tph2 was significantly reduced in the morphant and rescued by mRNA overexpression. Bb–c: The levels for tph1a and tph1b were unchanged. C: Serotonin immunoreactivity was reduced in the morphant. Ca: Std ctrl. Cb: Ntrk2b MO. Cc: Ntrk2b+mRNA. D: Expression of serta was observed in the raphe and pretectal cluster in Da (Std ctrl), Db (Ntrk2b MO), and Dc (Ntrk2b+mRNA). The expression was reduced in the morphant Db. Partial rescue was observed with mRNA overexpression in the morphants Dc. E: The Ntrk2b knockout mutants had reduced serotonin immunoreactivity as compared to wildtype littermates. At 6 dpf, Ea (wildtype) and Eb (homozygote mutant) are represented with reduced immunoreactivity in the hypothalamus. Ec–d. Adult sections from wildtype and knockout brains samples in the hypothalamic population. The expression of 5-HT in the mutant is aberrant. PVOa – paraventricular organ, anterior part, PVOi – paraventricular organ, intermediate part, PVOp – paraventricular organ. * p =0.0252 one-way ANOVA.Scale bar =100 µm.

In the Ntrk2b mutant, 5HT immunoreactivity was assessed in both the larval brain and adults. We observed a reduction in all the 5HT cell groups in the whole brain of the larval mutant at 5 dpf (Fig. 4Ea–b). No cell group was found to be completely abolished. In the Ntrk2b mutant adult brain sections, at a higher magnification of 5HT, immunoreactivity in the hypothalamic cell groups exemplified reduced neuronal cells in two different serotonergic hypothalamic neuron populations called the periventricular organ intermediate part (PVOi), close to the midline in the posterior tuberculum, and the periventricular organ posterior (PVOp) in the ventrocaudal hypothalamus (Fig. 4Ec–d). The effects of Ntrk2b loss were seen in the overall serotonergic cell populations.

### Consequences of using the trk inhibitor K252a on larval and adult zebrafish

The tyrosine kinase inhibitor K252a has commonly been used to study the inhibition of Trk signaling. To examine the effect of K252a on zebrafish, we analyzed two different concentrations in larval zebrafish at 5 dpf, when no major effect on fish mortality was observed. Western blotting with the pTrkB antibody was performed to determine the effective dose. TrkB phosphorylation was eliminated at 200 nM using an antibody specific for phosphorylated TrkB (pTrkB) (Fig. 5Aa). The total Trk levels measured using a polyclonal Trk antibody (Sc-11) remained unchanged (Fig. 5Ab). The loading control used in this study was actin and the levels were unaltered (Fig. 5Ac). The effective concentration for this study were about 100 nM and 200 nM, with no significant effect on fish swimming behavior. In the 5 dpf larvae, TH1 and 5HT immunoreactivity was analyzed in the treated fish (Fig. 5Ba–f). With K252a treatment at 200 nM, serotonin immunoreactivity was completely abolished with very few cells visible in the hypothalamus (Fig. 5Bc). TH immunoreactivity in these fish was reduced at both concentrations of K252a, 100 nM and 20 nM (Fig. 5Bd–f). These results were quite consistent with the morpholino as well as the mutant experiments.

**Fig 5.**
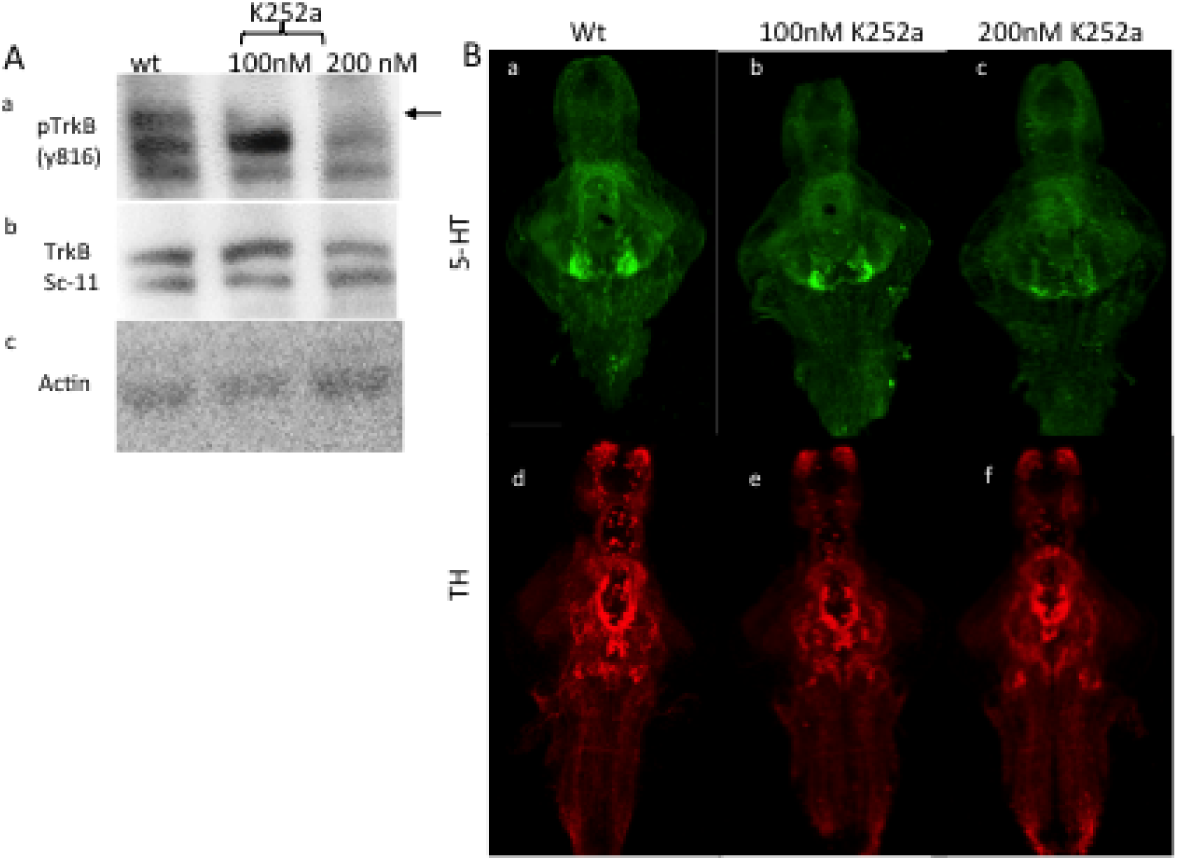
Pharmacological effects of K252a on TrkB phosphorylation and disruptions in aminergic populations. Aa: pTrkB antibody detected a phosphorylation band in the wildtype which dose-dependently disappeared in the 200 nM-treated adult fish brain samples. Ab: The total Trk bands were unaffected. Ac: Actin was used as the loading control. B (a–c): 5-HT immunoreactivity (ir) was detected in the wildtype untreated fish; 100 nM has fewer cells detected in the hypothalamus and the ir was completely abolished at 200 nM. B (d–f): TH ir in the diencephalon was disturbed. Scale bar = 100 µm.

## Discussion

In the present study, we established an Ntrk2b knockout mutant and demonstrated a crucial role of this receptor in the maintenance of the aminergic system in the brain. The *ntrk2b* loss-of-function studies during development were carried out using translation-blocking morpholino oligonucleotides (MO). No gross phenotype was observed in the MO-injected fish. Due to the limitations of MO methods (Schulte-Merker and Stainier, 2014), we also investigated the effects using the null mutant for comprehensive analysis of *ntrk2b* function. The validity of the results was verified using appropriate controls.

The main function of *ntrk2b* we observed was the maintenance of the two important aminergic systems involved in stress, depression, and antidepressant-mediated responses: the dopaminergic and the serotonergic system. In the Ntrk2b-deficient models, the dopaminergic cell population found to be affected was the ventral diencephalic cluster, which corresponds to the mid-brain dopaminergic cell groups A9/A10 in mammals (Rink and Wullimann, 2002). Brain-derived neurotrophic factor (BDNF) is known to exert its trophic function on the mesencephalic dopaminergic neurons (Hyman et al., 1991). These neuronal populations have been linked with several neurological disorders such as Parkinson’s disease, schizophrenia, addiction, and psychomotor retardation. *In vitro* studies on mesencephalic dopamine neurons suggest that the effect of BDNF via TrkB receptor activation is responsible for dopamine release (Blochl and Sirrenberg, 1996). The absence of the Ntrk2b receptor abolished BDNF function in a subset of the dopaminergic neuronal population in the zebrafish, which might be essential for depressive-like behavior and may also be responsible for the antidepressant response efficacy. Further studies regarding depressive behavior using these mutants could possibly confirm the regulatory role of Ntrk2b on these dopaminergic neurons.

BDNF and its high-affinity cognate receptor, tyrosine kinase B (TrkB), have been concomitantly related to serotonin (5-HT) in a myriad of neurochemical and behavioral responses following SSRI treatment. Therefore, we investigated the serotonergic neuronal cell groups in these mutants. Serotonin cell populations have been very robustly studied and characterized in the zebrafish (Lillesaar et al., 2009; Sallinen et al., 2009). Unlike dopamine circuitry, all the different serotonin populations were reduced. The anterior ascending axonal projections were found to be disrupted. The markers for serotonin synthesis and re-uptake, *tph2* and *serta*, were significantly reduced, suggesting a direct role of *ntrk2b* in serotonergic neuron development and maintenance. These results suggest some important regulatory mechanism between the BDNF/TrkB and serotonin. Thus, these mutants could potentially be used to dissect the complex neuronal circuitry associated with mood-related behaviors.

Zebrafish are amenable to drug screens because the larvae or adults can be exposed to the drugs by the addition of the chemicals to the swimming tank water. The *ntrk2b* mutants in the future could be useful for small molecule screening and could potentially be used in antidepressant drug discovery. The sequence homology for *ntrk2b* in zebrafish with that of humans and mice is 62%. The zebrafish retains about 20% of duplicated gene pairs. The pleiotropism of each duplicated gene at a single gene locus that has a cause or an effect on similar physiological processes is therefore reduced. This, in turn, facilitates genetic study in these species. The complete knockout mutants for both BDNF and TrkB in rodents and mammals die due to peripheral abnormalities. This has been a limitation in understanding the cause or effect in the signaling pathways during development. Interestingly, the viability of the Ntrk2b zebrafish knockout could probably be due to sub-functionalization in the cis-regulatory elements of the duplicated genes, or then an adaptive modification of the duplicated gene clusters (Prince and Pickett, 2002). Thus, zebrafish could serve a good model to understand the biochemical and physiological analysis of the Ntrk2 function. This offers promising avenues to decipher the physiological processes underlying psychiatric disorders involving BDNF/TRKB signaling.

## Acknowledgements

We thank Outi Nikkila, Sulo Kohlemainen, Riikka Pesonen, and Reeta Huhtala for their expert technical help, and Henri Koivula for the fish husbandry and sampling. This work was financially supported by ERC grant No 322742 – iPLASTICITY, the Sigrid Juselius Foundation, and Academy of Finland grants #294710 and #307416 to EC

